# FAF2 is a bifunctional regulator of peroxisomal homeostasis and saturated lipid responses

**DOI:** 10.1101/2024.12.12.628015

**Authors:** Choah Kim, Katlyn R. Gabriel, Dylan Boone, Matthew R. Brown, Katherine Oppenheimer, Maria Kost-Alimova, Juan Lorenzo B. Pablo, Anna Greka

## Abstract

Exposure to saturated fatty acids (SFAs), such as palmitic acid, can lead to cellular metabolic dysfunction known as lipotoxicity. Although canonical adaptive metabolic processes like lipid storage or desaturation are known cellular responses to saturated fat exposure, the link between SFA metabolism and organellar biology remains an area of active inquiry. We performed a genome-wide CRISPR knockout screen in human epithelial cells to identify modulators of SFA toxicity. The screen revealed peroxisomal proteins, especially those that impact ether lipid synthesis, as important regulators of lipotoxicity. We identified Fas-associated factor family member 2 (FAF2) as a critical bifunctional co-regulator of peroxisomal and fatty acid biology. We further uncovered a new biological function for the ubiquitin-regulatory X (UBX) and UAS thioredoxin-like domains of FAF2, demonstrating their requirement for peroxisomal protein abundance and SFA-induced cellular stress. Our work highlights the role of FAF2 in regulating peroxisomal abundance and function, and the peroxisome as a key organelle in the cellular response to SFAs.

## Introduction

Fatty acids are important metabolic molecules, serving as energy reservoirs, building blocks of cellular membranes, lipid signaling molecules, and fuel for energy generation. Various adaptive processes exist to help maintain fatty acid homeostasis, including their storage in lipid droplets, incorporation into membrane lipids, use as signaling molecules, and breakdown through beta-oxidation in mitochondria or peroxisomes. Disruption of these adaptive processes results in cellular injury, and specifically, prolonged exposure to excess free fatty acids (FFAs) induces lipotoxicity, characterized by activation of the unfolded protein response (UPR), oxidative stress, inflammation, and apoptotic or ferroptotic cell death (*1–3*). Genome-wide screens in human Jurkat and K562 cells have revealed key mechanisms of saturated fatty acid-mediated lipotoxicity (*4*, *5*). However, we recently demonstrated that diverse free fatty acid species can drive unique cellular responses in a cell-type specific manner (*6*). As such, there remains a significant and fundamental knowledge gap in our systematic understanding of lipotoxic responses in diverse human cell types.

Lipid homeostasis is maintained through the compartmentalization of metabolic processes to specialized organelles such as peroxisomes (*7*, *8*). Peroxisomes are well known for their ability to metabolize reactive oxygen species (ROS), breakdown fatty acids through beta-oxidation by collaborating with the mitochondria, and synthesize ether lipids (*7*, *9–11*). Both the ER and mitochondria are required for the assembly of peroxisomes by supporting the transport of peroxisomal biogenesis factors, or peroxins (PEX proteins), to new peroxisomal membranes (*12*, *13*). Mutations in the 12 known human PEX genes disrupt organelle assembly or function and lead to peroxisome biogenesis disorders (*14–16*). Peroxisomal biogenesis involves the sequential incorporation of peroxisomal membrane proteins (PMPs) into pre-peroxisomal vesicles followed by the import of matrix proteins that are important for peroxisomal functions. Peroxisomal matrix proteins such as mevalonate kinase (MVK), alkylglycerone phosphate synthase (AGPS), and glyceronephosphate O-acyltransferase (GNPAT), contribute to the biosynthesis of cholesterol and ether lipids (*11*, *17*, *18*). Peroxisomal integrity is therefore critical for the diverse metabolic roles of peroxisomes, but bifunctional regulators of peroxisomal homeostasis and the lipotoxicity response are still largely unexplored.

Here, we used CRISPR-based genome-wide screening of human epithelial cells to systematically explore mechanisms of cellular lipotoxicity. We integrated our results with publicly available datasets to identify both shared and cell-specific pathways that determine cellular fitness after exposure to SFAs. We found that depletion of peroxisomal proteins protects cells from palmitic acid-induced lipotoxicity and that Fas associated factor 2 (FAF2) regulates peroxisomal abundance. Our work reveals a general, previously unrecognized bifunctional role for FAF2 in regulating peroxisomes and cellular responses to lipotoxic saturated fatty acids.

## Results

### Genome-wide screen identifies a unique peroxisomal signature in the epithelial cell response to palmitic acid

Palmitic acid (PA) is the most prominent saturated fatty acid (SFA) in human serum and increased levels are correlated with the development of a multitude of metabolic disorders that affect different organ systems and tissues (*19–21*). Currently there is a limited understanding of the mechanisms regulating lipid-induced injury and how they differ across diverse cell types. Given recent interest in SFA-mediated injury in kidney, lung, and liver epithelia leading to fibrotic diseases (*22–24*), we explored mediators of PA-induced cellular toxicity through a genome-wide loss-of-function viability screen in human epithelial cells. We infected a library of approximately 80,000 guide-RNAs (gRNAs) targeting ∼20,000 genes into immortalized human epithelial cells derived from a normal human kidney (*25*, *26*). After puromycin selection, we treated cells with either the fatty acid carrier, BSA, or BSA-conjugated PA (*6*). After 14 days, we harvested and sequenced pooled cells, and analyzed for enrichment or depletion of guides compared to non-targeting or no-gene controls (Figure 1A). A doubling time of ∼2 days was maintained with BSA conditions, while PA treated cells exhibited a growth disadvantage with a doubling time of ∼3.2 days (Figure S1A). The screen was conducted in duplicate, showing robust discovery of established lipotoxicity modulators such as Acetyl-CoA Acetyltransferase 2 (ACAT2), Calcineurin Like EF-Hand Protein 1 (CHP1), and Acyl-CoA Synthetase Long Chain Family Member 3 (ACSL3), as well as newly identified mediators such as Fas-associated factor 2 (FAF2) (Figure 1B).

**Figure 1:**
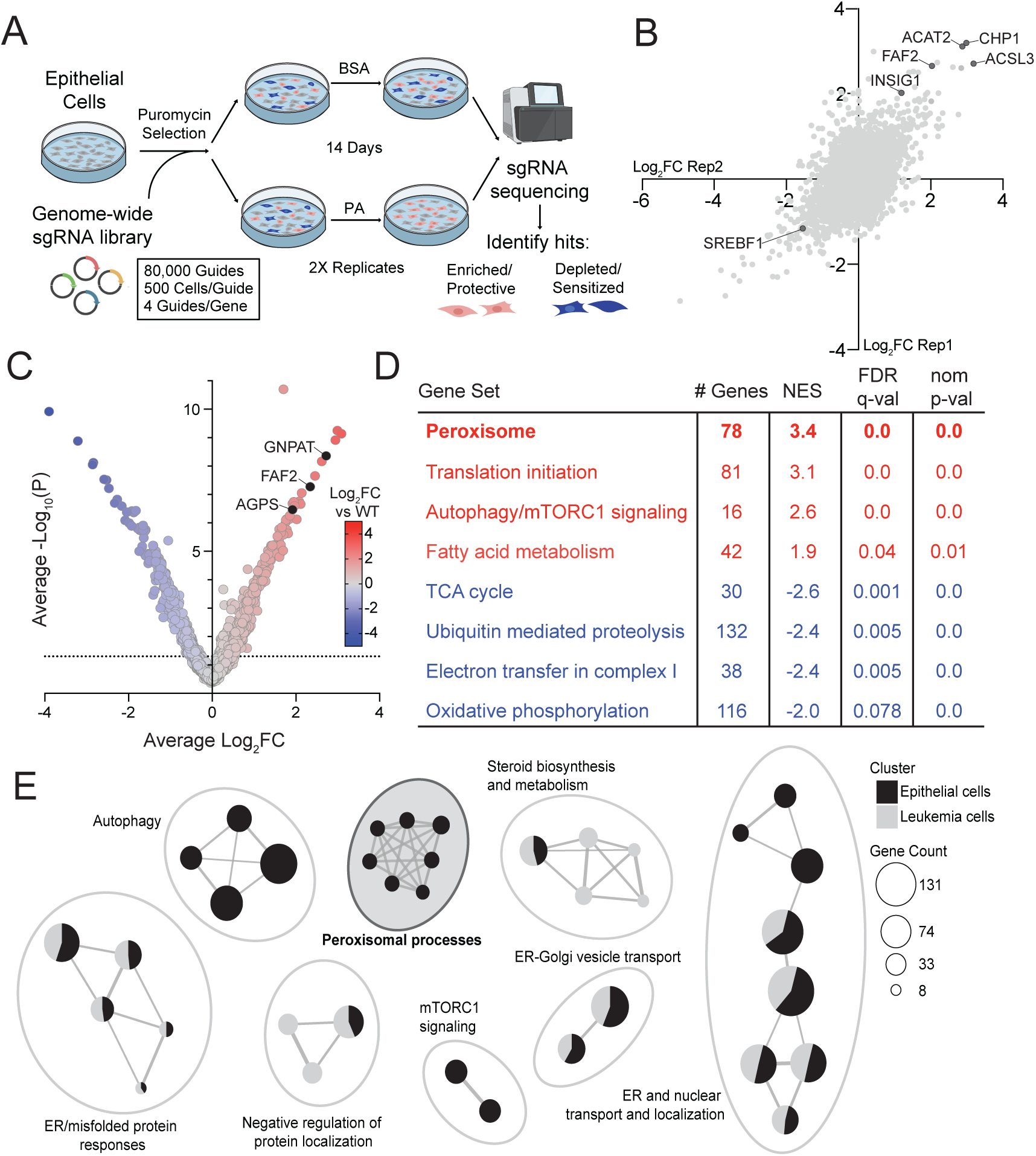
Genome-wide screen reveals peroxisomal pathways as mediators of lipotoxicity. (**A**) Experimental design of the genome-wide CRISPR/Cas9 knockout viability screen. Cells were treated with BSA or 500 µM PA for 14 days to identify genetic knockouts that sensitize or protect cells from PA-induced toxicity (P<0.05). (**B**) Correlation plot showing reproducibility of screen hits among replicates 1 and 2 of the CRISPR knockout viability screen treated with BSA or 500 µM PA. (**C**) Volcano plot of screen hits in epithelial cells. All genes were plotted for their Log2-fold change (Log2FC) enrichment in PA versus BSA conditions and significance level (-Log10(P)). (**D**) Gene set enrichment analysis (GSEA) of PA screen hits showing select KEGG pathways that are most significantly enriched or depleted in epithelial cells. FDR<25%. (**E**) Enrichment analysis of GO terms comparing significant hits from PA screens in epithelial cells or K562 leukemia cells identifies unique lipotoxicity modulators between cell types. The clustered network plot was generated using clusterProfiler with p-adjusted value cutoff adjP<0.25. Correction for multiple testing was performed using the Benjamini-Hochberg method. The gene count of a node is the number of genes within each GO term.

We identified 966 genes that, upon deletion, protected cells from PA-induced lipotoxicity and 847 genes that were knockout (KO)-sensitizing (P<0.05) (Figure 1C). Gene set enrichment analysis (GSEA) of our dataset revealed positively enriched KEGG pathways, such as peroxisome, autophagy/mTORC1 signaling, and fatty acid metabolism, while various mitochondrial processes like the electron transport chain and oxidative phosphorylation complex I, as well as ubiquitin-mediated proteolysis, contributed to the most sensitizing lipotoxicity modulators (Figure 1D). Following discovery of enriched pathways in epithelial cells, we then compared our results with a previously published screen in leukemia cells (*4*) (Figure S1B). This integrated analysis of two independent lipotoxicity screens allowed for the identification of both unique and common modulators of lipotoxicity (Table S1). First, we performed DAVID functional annotation clustering on the 99 genes that were common hits between the two screens to identify significant biological pathways emerging from this shared gene set (Figure S1C). Sterol synthesis and fatty acid metabolism were among the top enriched processes. Consistent across both cell types, knockout of the SREBF1 pathway genes (SREBF1 and INSIG1) greatly modified the cells’ ability to respond to PA treatment (Figure 1B and Table S1). Clustered network analysis also revealed shared enriched processes, such as regulation of ER protein localization, sterol biosynthesis, and misfolded protein responses. Intriguingly, we identified a cluster of peroxisomal gene sets that was strongly enriched and specific to epithelial cells (Figure 1E).

### Systematic data integration reveals FAF2 as a key regulator of epithelial cell lipotoxicity and peroxisomal biology

Having identified peroxisomes as significant regulators of the response to PA-induced lipotoxicity in epithelial cells, we sought to identify the organellar processes that contribute to this phenotype. Expanding our GSEA search to a broader set of gene set databases, including KEGG, GOBP, and REACTOME, we found that many peroxisomal processes, including ether lipid metabolism, bile acid metabolism, and peroxisomal protein import and transport, contribute to PA-induced toxicity in epithelial cells (Figure 2A, Figure S2A). Further, the guides that contributed to this peroxisomal gene set were uniquely enriched in epithelial cells (Figure 2B). Of genes with annotated peroxisomal functions, ether lipid synthesis enzymes were distinctly the most significantly KO-protective in our screen, while peroxisomal matrix proteins that are involved in other lipid metabolic processes, like beta-oxidation enzymes, were not significant (Figure 2C).

**Figure 2:**
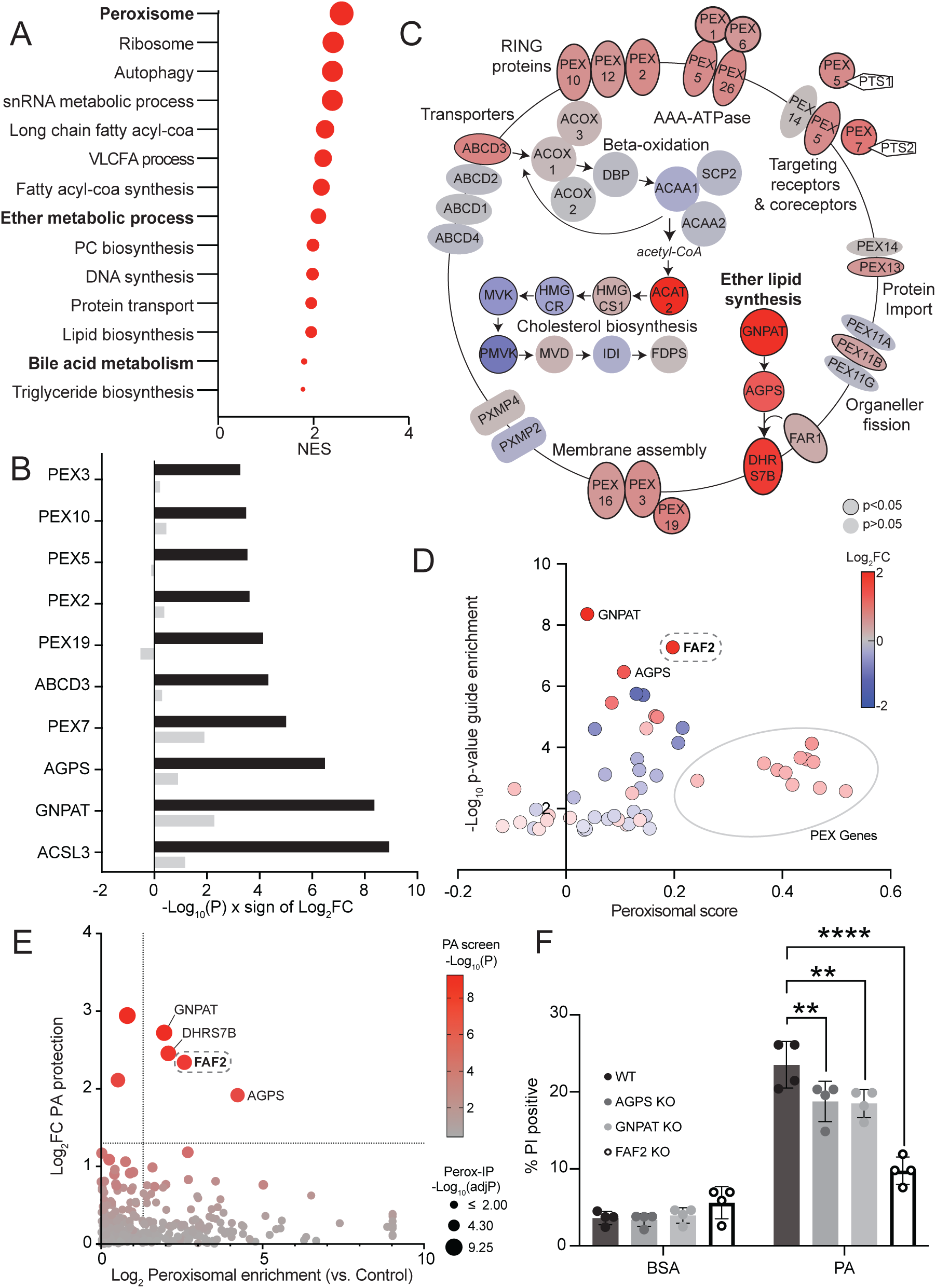
FAF2 is a putative bifunctional regulator of peroxisomes and cellular responses to lipotoxicity. (**A**) GSEA analysis of gene knockouts that enrich or deplete in PA conditions shows high enrichment of peroxisomal processes. GSEA normalized enrichment scores (NES) are displayed for differentially enriched sets of guides; gene sets include KEGG, GOBP, and REACTOME pathway entries. The size of the data points corresponds to the NES. (**B**) Top-ranking enriched guides that contributed to the KEGG peroxisomal gene set are significantly KO-protective in epithelial cells (black) but not K562 leukemia cells (gray). (**C**) Peroxisomal genes are organized by their functional process with red indicating genes that are KO protective and blue indicating genes that are KO sensitizing. Genes with a black outline indicate P<0.05 in the PA screen. (**D**) FAF2 and ether lipid synthesis enzymes lie at the intersection of lipid and peroxisomal biology. Genes are plotted by coessentiality scores with peroxisomal biogenesis factors (x-axis) and enrichment in the PA screen (y-axis). Correlation with peroxisomal biogenesis factors was calculated as a peroxisomal score, the average Pearson correlation score of each gene vs PEX genes (DepMAP CRISPR, Chronos). (**E**) The Log2 enrichment (y-axis) of all genes that were significantly enriched in the PA screen plotted against enrichment in cellular peroxisomal IP fractions (x-axis). Multiple unpaired t-tests with two-stage step up (Benjamini correction FDR=1%) was calculated for peroxisomal enriched samples compared to control IP. Ether lipid synthesis genes and FAF2 are enriched in both datasets. (**F**) Loss of peroxisomal ether lipid synthesis enzymes or FAF2 protects cells from PA-induced toxicity over a five day growth period. Cells were treated with BSA or 125 uM PA for five days and measured for cell death using propidium iodide (PI) dye. ****p<0.0001. Ordinary two-way ANOVA with Tukey’s multiple comparisons test, with a single pooled variance.

Annotation of peroxisomal or ether lipid synthesis pathways are historically lacking in mammalian systems and are still being actively investigated. Recent progress has been made in discovering novel functions of poorly characterized genes with the use of computational approaches such as coessentiality mapping (*27*). We therefore sought to identify bifunctional modulators of peroxisomal processes and lipotoxicity by combining coessentiality mapping with genes identified as lipotoxicity modulators from our screen. To do this, we generated a peroxisomal functional score for each gene hit from our PA screen by averaging the Pearson correlation scores from DepMap (a database derived from ∼1000 cancer cell lines (*28*)) between that gene and a curated list of every peroxin (PEX) gene important for the formation and function of peroxisomes. Plotting the peroxisomal score against the significance of enrichment or depletion in our screen highlighted genes likely to be important for both peroxisomal function and the cellular response to PA. Among our PA screen hits, we identified a list of 52 genes functionally related to peroxisomal biogenesis factors among which we searched for new candidate regulators of peroxisomes. Foremost among these genes were known peroxisomal and lipid biology genes, such as peroxisomal ether lipid synthesis enzymes AGPS and GNPAT (*11*), in addition to highly scoring PEX genes that served as positive controls (Figure 2D). Most importantly, this analysis revealed FAF2 as a gene without any prior peroxisomal functional annotations. The emergence of FAF2 from an analysis that integrated DepMap with our epithelial cell CRISPR screen results implied that FAF2 may serve as a general regulator of peroxisomes across many human cell types (Figure 2D and S2B-D).

In a parallel approach aimed at exploring possible direct regulators of peroxisomes, we searched for screen hits that localize to peroxisomes. We utilized a dataset that identified proteins enriched by organellar isolation in T47-D human epithelial cells to identify PA screen hits that may localize to peroxisomes (*29*). Ether lipid synthesis proteins AGPS, GNPAT, and DHRS7B were all significantly enriched in peroxisomal IP fractions, in addition to being protective hits in our screen (Figure 2E). Among novel regulators, we again identified FAF2 as a top hit that was significantly enriched in peroxisomal IP fractions (Figure 2E). Having functionally mapped FAF2 as a putative peroxisomal regulator, we assessed how cells respond to loss of FAF2 or peroxisomal ether lipid synthesis enzymes, AGPS or GNPAT. In agreement with the results of our screen, genetic knockout of FAF2, AGPS, and GNPAT, protected from cell death after five days of PA exposure (Figure 2F). Thus, FAF2 emerged as a new putative bifunctional regulator of peroxisomes and of PA-induced cellular toxicity.

### FAF2 is generally required for peroxisomal protein abundance and ether lipid synthesis

To explore the mechanisms through which FAF2 and the ether lipid synthesis enzymes AGPS and GNPAT regulate peroxisomal and saturated lipid biology, we used tandem mass tag liquid chromatography-mass spectrometry (TMT LC-MS) to quantify proteomic changes in FAF2, AGPS, and GNPAT KO cells (Figure 3A, Figure S3A-B, Table S2). We hierarchically clustered proteomic level changes in these three KO lines and found that peroxisome proteins, such as PEX1 and PEX3, were consistently reduced (Figure 3B). These data were consistent with a putative role for FAF2 in the regulation of peroxisomes (Figure 2) and further suggested that the ether lipid synthesis enzymes AGPS and GNPAT may also be important for maintaining peroxisomal protein abundance.

**Figure 3:**
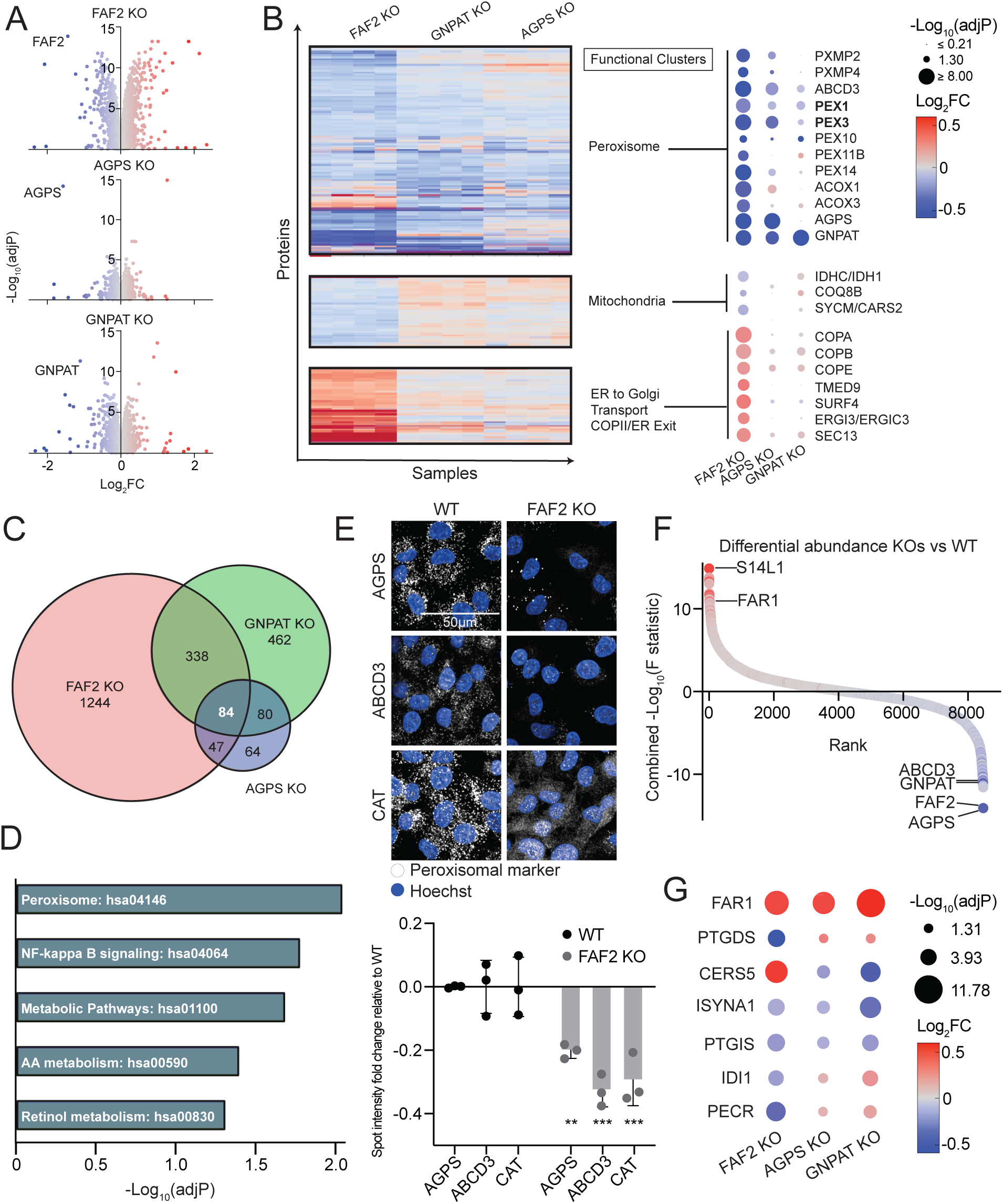
FAF2 KO leads to changes in the abundance of peroxisomal proteins and proteins that regulate lipids. (**A**) Volcano plots of Log2 fold change (x-axis) plotted against -Log10(adjP) significance (y-axis) of normalized proteins in FAF2 KO, AGPS KO, and GNPAT KO cells compared to wildtype cells. (**B**) Hierarchical clustering of proteomics revealed 3 distinct clusters of proteins commonly changed between all KO cell lines. Log2 fold change (Log2FC) was calculated for normalized protein intensity values of knockout samples versus the average of wildtype cells, with proteins increased in KO samples (in red) or decreased (in blue). (**C**) 84 proteins with significantly altered abundance in all three KO cell lines were identified. (**D**) DAVID GO functional clustering highlights overrepresented processes that are commonly enriched or depleted across all three KO cell lines. We conducted functional enrichment of biological processes of proteins belonging to the clusters identified in Fig 3B. Adjusted Benjamini values (-Log10(adjP)) are reported for discovered processes. (**E**) Immunofluorescence imaging of peroxisomal proteins, AGPS, ABCD3, and catalase (CAT) in wildtype (WT) and FAF2 KO cells. ***p<0.001. Ordinary two-way ANOVA with Šídák’s multiple comparisons test. Scale bar = 50 µm. (**F**) The mean difference between experimental groups and wild-type was calculated to determine proteins that are changed in the KO cell lines. Genetic disruption resulted in decreased abundance of the proteins encoded by the corresponding targeted genes (FAF2, GNPAT, AGPS) as well as some commonly altered proteins (FAR1, S14L1, and ABCD3). (**G**) Commonly altered protein levels in KO cell lines. Significantly increased proteins are shown in red while significantly decreased proteins are shown in blue (size indicates -Log10(adjP)).

FAF2 KO cells showed the most notable shift in the total proteome (i.e. number of protein abundances changed and magnitude of change). This observation was in line with the previously described role of FAF2 in proteasome-dependent degradation (*30–32*), and accordingly, known FAF2 targets were increased in abundance in FAF2 KO cells (Figure S3C). Of note, FAF2 KO also resulted in the largest disruption of peroxisomal protein abundance: it reduced both peroxisomal matrix (AGPS, GNPAT, ACOX1/3) and peroxisomal membrane proteins (PXMP2/4, ABCD3), demonstrating a critical role for FAF2 in maintaining peroxisomal protein abundance upstream of ether lipid synthesis enzymes (Figure 3B). FAF2 KO also resulted in decreased mitochondrial protein levels and increased abundance of ER transport and proteostasis proteins (Figure 3B). Importantly, FAF2 emerged as a general, conserved regulator of peroxisomal protein abundance because peroxisomal and mitochondrial proteins were also decreased in a publicly available proteome of FAF2 KO in HEK293T cells (Figure S3D-E) (*33*). Since we did not observe significant changes in mRNA levels of most peroxisomal genes (Figure S3F), we concluded that FAF2 likely regulates peroxisomal abundance post-transcriptionally.

There were 84 proteins whose abundance was changed across all three KO lines (Figure 3C). Pathway analysis of these 84 proteins using DAVID GO confirmed robust changes in peroxisomal pathways and identified changes in metabolic and inflammatory pathways (Figure 3D). Immunofluorescence imaging corroborated that FAF2 KO leads to loss of peroxisomal proteins AGPS and ABCD3 (Figure 3E). We also observed that catalase (CAT), one of the most abundant proteins in peroxisomes (*9*) was no longer concentrated in its characteristic punctate pattern – instead, CAT showed a diffuse cytosolic signal consistent with peroxisomal disruption (Figure 3E) (*34*, *35*). Depletion of cellular levels of peroxisomal proteins ABCD3, GNPAT, AGPS, PEX5, and PEX3 (but not of the soluble protein PEX19 or matrix enzyme CAT) was also observed by western blot in FAF2 KO cells (Figure S3B). We predicted that in response to the genetic disruption of ether lipid enzymes, there would be a compensatory increase in FAR1, whose activity is directly inhibited by ether lipids (*36*), which was indeed validated across each of the KO lines (Figure 3F-G, 4A). Additionally, we found that several metabolic or lipid enzymatic proteins were differentially expressed, leading us to hypothesize that FAF2, GNPAT and AGPS KO cells have an altered lipid metabolic state (Figure 3G).

### Peroxisomal ether lipids regulate the cellular response to saturated fatty acids

The enzymes GNPAT and AGPS initiate ether lipid synthesis in the peroxisomal matrix, which is then completed in the endoplasmic reticulum (Figure 4A) (*37*, *38*). Our proteomics results indicated that FAF2 functions upstream of GNPAT and AGPS because FAF2 KO reduced the abundance of GNPAT and AGPS. We thus assessed changes in the lipidomes of cells depleted of AGPS, GNPAT, and FAF2, as well as PEX3 and PEX5, additional proteins upstream of ether lipid synthesis important for peroxisomal membrane and matrix protein import, respectively (*39*, *40*). Ether lipid species, plasmenyl PEs and PCs, were the most consistently depleted class of lipids across all five genetic knockouts demonstrating the importance of peroxisomal integrity to ether lipid synthesis (Figure 4B-C, S4A, S3B, and Table S3). The decrease of plasmenyl lipid species in FAF2 KO cells also lent support to the role of FAF2 in peroxisomal function (Figure 4B-C and S4A). Further, the decrease in plasmenyl lipids in the knockout lines could be rescued by treatment with a post-peroxisomal ether lipid intermediate, 1-O-hexadecyl-sn-glycerol (EL), but not by the addition of an ester-linked equivalent (ES) (Figure 4D).

**Figure 4:**
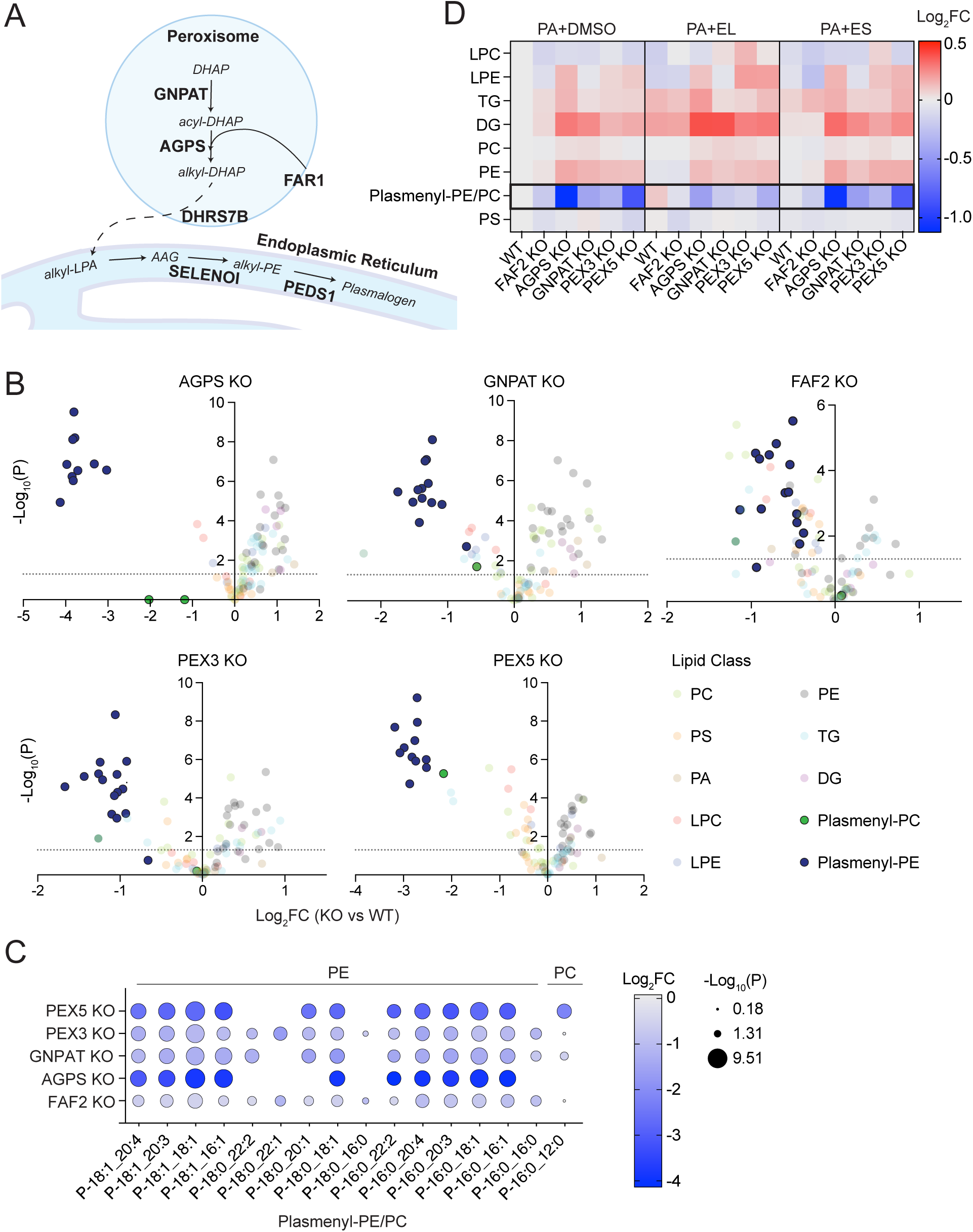
Disruption of peroxisomes leads to ether lipid depletion and protects cells from saturated fatty acid exposure. (**A**) Schematic representing essential genes in the ether lipid synthesis pathway. (**B**) Volcano plots showing Log2 fold change of normalized lipid intensities in peroxisomal knockout cells show a decrease in plasmenyl lipid species compared to wildtype cells treated with PA. Dotted line shows cutoff for lipid fold changes that are significant at p<0.05. (PC=phosphatidylcholine; PS=phosphatidylserine; PA=phosphatidic acid; LPC=lysophosphatidylcholine; LPE=lysophosphatidylethanolamine; PE=phosphatidylethanolamine; TG=triacylglycerol; DG=diacylglycerol; Plasmenyl-PC=plasmenyl-phosphatidylcholine; Plasmenyl-PE=plasmenyl-phosphatidylethanolamine). (**C**) Plasmenyl-PEs and PCs are globally downregulated in PA-treated FAF2 and peroxisomal knockout cells compared to wildtype cells. (**D**) Heatmap of various lipid species’ Log2 fold changes (Log2FC) across genotypes showing a decrease in ether lipid species in peroxisomal knockout cells, which is reversible with ether lipid treatment, but not ester lipid. Cells were treated with PA+DMSO, PA+ether lipid (EL) intermediate (1-O-hexadecyl-sn-glycerol), or PA+ester lipid (ES) (dl-a-palmitin) as a control.

We also observed an increase in diacylglycerol (DG) and triacylglycerol (TG) levels with PA treatment in the peroxisomal and FAF2 KO lines compared to control cells (Figure 4D, Figure S4A). Cells can deal with excess FFAs by incorporating them into TGs, which are stored in lipid droplets (LDs) (*41*). Since previous work has shown that FAF2 regulates LD degradation through Adipose Triglyceride Lipase (ATGL) (*42*), we sought to assess whether an increase in LDs could explain the protective effect against PA toxicity in FAF2 KO cells. To this end, we tested whether inhibition of lipid droplet formation using Diglyceride O-acyltransferase (DGAT)1 or DGAT2 inhibitors could reverse protection against PA. We found that cell viability was not affected even in cells depleted of lipid droplets (Figure S4B).

In addition to affecting LDs, PA is known to induce ER stress and the unfolded protein response (UPR) (*2*, *4*). Therefore, we measured UPR markers ATF4 and XBP1s and found that FAF2 KO was protective: it decreased ER stress caused by PA treatment (Figure S4C). The protective effect was reversed by addition of ether lipids (i.e. EL), suggesting that the presence of ether lipids is sufficient to promote the toxic effects of PA. Taken together, these studies showed that the PA-protective effect in FAF2, AGPS and GNPAT KO cells (Figure 2F) is through disruption of peroxisomal ether lipid synthesis and not due to an increase in lipid storage in LDs (Figure S4B).

### The UBX and UAS domains of FAF2 regulate peroxisomal abundance and the response to saturated fatty acids

FAF2 is composed of three major domains, ubiquitin-associated (UBA), UAS thioredoxin-like, and the ubiquitin-binding X (UBX) domains (Figure 5A) (*43*). The UBA domain is known to recognize ubiquitin-tagged substrates, the UAS domain interacts with unsaturated long chain fatty acids, and the UBX domain interacts with p97/VCP to regulate ER-associated degradation (ERAD) (*31*, *42–46*) – however, their role in peroxisomal lipid biology remains unknown. We tested whether a specific domain in FAF2 is responsible for peroxisomal regulation by expressing a panel of domain-deleted constructs in FAF2 KO cells (Figure 5A-B). As expected, FAF2 KO reduced protein levels of both ether lipid enzymes, AGPS and GNPAT, as well as PEX10 as shown by western blot (Figure 5C). Expressing full length FAF2 (FAF2 full) or FAF2 without the UBA domain (FAF2ΔUBA) in FAF2 KO cells restored the abundance of these peroxisomal proteins. In contrast, expression of FAF2 lacking the UBX (FAF2ΔUBX) or UAS (FAF2ΔUAS) domains led to only a partial restoration of these proteins (Figure 5C). Similarly, the dispersion of CAT and loss of AGPS in FAF2 KO cells was partially reversed to peroxisome-like punctae by FAF2 full and FAF2ΔUBA but not FAF2ΔUBX or FAF2ΔUAS expression (Figure 5D). Total CAT protein levels did not change (Figure S5). Consistent with the hypothesis that intact peroxisomal morphology and function are necessary for ether lipid-mediated PA toxicity, FAF2ΔUBX or FAF2ΔUAS protected cells from PA, while introducing FAF2 full or FAF2ΔUBA reversed the protective effect of FAF2 KO as measured by propidium iodide positive nuclei (Figure 5E). We concluded that the UBX and UAS domains of FAF2 were necessary both for peroxisomal abundance and for the cellular response to PA.

**Figure 5:**
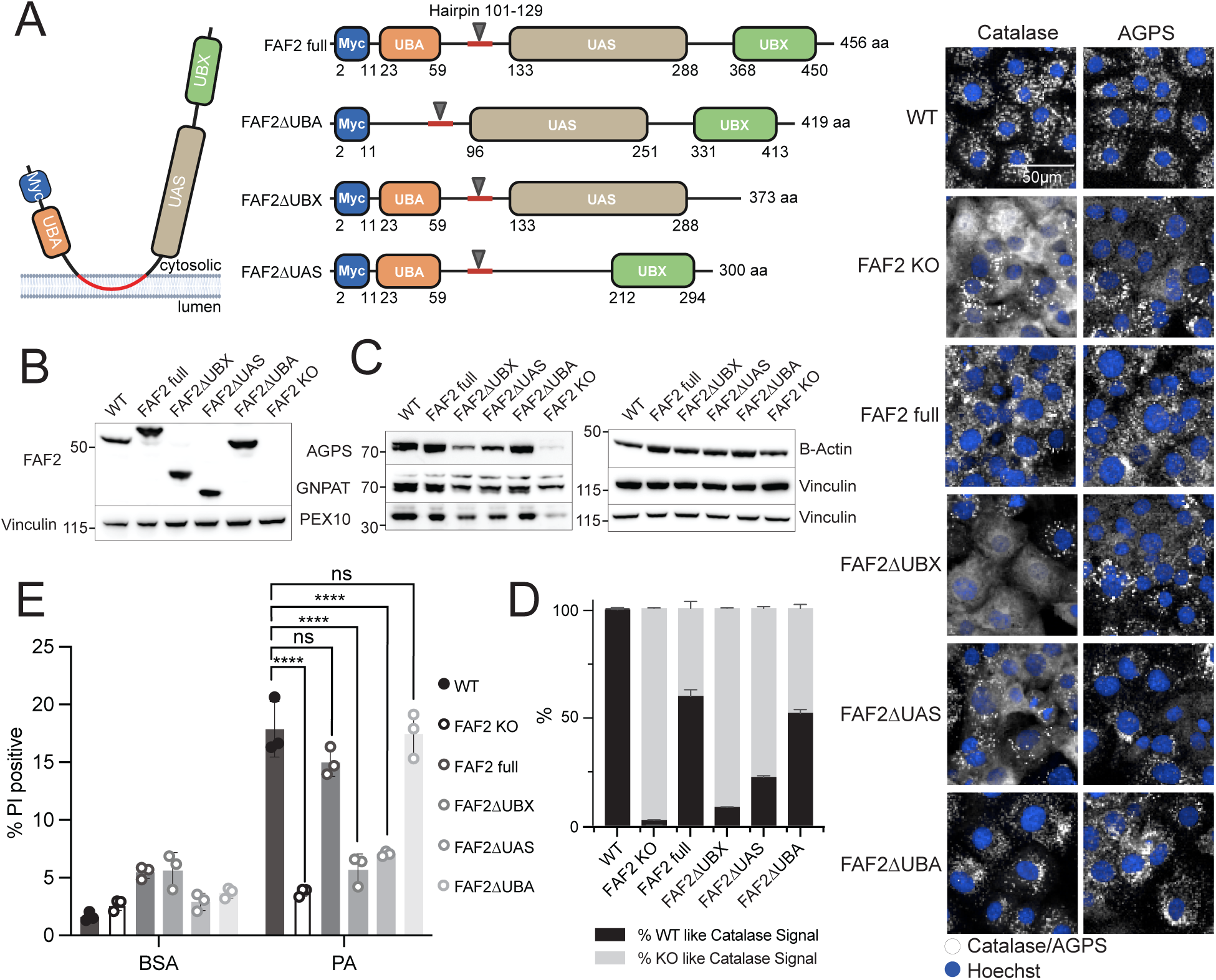
FAF2 mediates saturated fatty acid toxicity and peroxisomal abundance via its UBX and UAS domains. **(A)** Schematic of the FAF2/UBXD8 hairpin-anchored protein and FAF2 domain deletion constructs. **(B)** Western blot of FAF2 in WT or FAF2 KO cells transduced with full or domain-deleted (ΔUBX, ΔUAS, or ΔUBA domain) myc-tagged FAF2 constructs. **(C)** Western blots of peroxisomal proteins AGPS, GNPAT, and PEX10 in WT, KO, and KO-cells with re-expressed full or domain-deleted FAF2 constructs (ΔUBX, ΔUAS, or ΔUBA domain) show that UBX and UAS domains are necessary to restore peroxisomal protein abundance. **(D)** (Left) CAT localization was quantified by training the dataset with Harmony high-content imaging software on WT and FAF2 KO phenotypes. Cells in each sample were then assigned to either a WT or FAF2 KO phenotype and percentage of results are displayed. N=3 biological replicates per genotype, quantification is of one representative experiment. (Right) Immunofluorescence images (60X) show that CAT punctae and AGPS protein abundance are dependent on the FAF2 UBX and UAS domains. Scale bar = 50 µm. **(E)** PA-induced toxicity is dependent on the UBX and UAS domains of FAF2. Cell death was quantified by percentage of propidium iodide (PI) positive cells. ****p<0.0001. Ordinary two-way ANOVA with Tukey’s multiple comparisons test.

## Discussion

Metabolism and organellar biology are inextricably linked, with many metabolic pathways compartmentalized in membrane bound environments and conversely, many organellar functions regulated by the metabolic state of the cell. Our current investigations into the cellular response to the lipotoxic fatty acid palmitate have uncovered a critical role for peroxisomal ether lipid synthesis in mediating SFA-induced lipotoxicity. Furthermore, our work has revealed a previously unrecognized bifunctional role for FAF2 in both peroxisomal biology and the regulation of lipotoxic cellular responses. There are several key implications from our work.

First, our study identified FAF2 as a critical mediator of ether lipid synthesis. FAF2 KO depleted ether lipids via disruption of peroxisomal abundance and induced more general changes in the cellular lipidome. These results expand our understanding of the cellular roles of FAF2 (*30–32*, *47*). Our results further suggest that the UBX and UAS – but not the UBA – domains of FAF2 are required for peroxisomal abundance as well as for lipotoxicity induction. The UBA domain is annotated as a ubiquitin binding domain (*48*), suggesting that recognition of ubiquitinated proteins is not necessary for the newly uncovered role of FAF2 in the regulation of responses to SFAs. We speculate that FAF2-p97/VCP segregase activity mediated through the UBX domain may contribute to regulation of peroxisomal proteins at ER-mitochondria contact sites (*33*, *47*). Consistent with this notion, FAF2 is known to be primarily localized in the ER and is enriched at ER-mitochondria contact sites, where its insertion is dependent upon PEX3 and PEX19 (*49*). Our data suggest that there may be reciprocal regulation in that FAF2 itself regulates cellular levels of PEX proteins that are important for peroxisomal integrity and the import of peroxisomal matrix proteins such as CAT and AGPS. Future studies may focus further on the precise mechanisms by which FAF2 regulates peroxisomes.

Second, our data suggest that ether lipid synthesis in the presence of saturated toxic fatty acids is a major determinant of lipotoxicity in human epithelial cells, an effect that was not observed in previous genome-wide lipotoxicity studies in other cell types i.e. cancer cells (*4*, *5*, *50*). This epithelial cell injury mechanism is highly relevant for many fibrotic diseases which are in aggregate associated with more than 45% of deaths in the industrialized world (*51*). Interestingly, prior work has shown that polyunsaturated ether lipid peroxidation contributes to ferroptotic cell death (*52*, *53*). Together with our current study, this work implicates peroxisomal derived lipids in a wide diversity of stress responses that likely depend on cell type and lipid properties (i.e., saturation level, such as polyunsaturated versus saturated). Since cellular and organismal health is based on interactions between genes and the environment, it is indeed likely that different cell types and tissues differentially utilize lipids for energy, signaling, and storage. Our studies underscore that when it comes to organellar biology and cell metabolism, comparative studies in diverse cell types (e.g. epithelial cells versus Jurkat or K562 cells) can provide insights into both general (i.e. FAF2’s regulation of peroxisomes) and cell-type specific mechanisms (i.e. the role of ether lipids in the epithelial cell SFA response). It is therefore necessary to expand our investigations into a broader array of cell types to precisely determine to what extent cell identity determines important regulatory pathways in cellular lipotoxicity. Finally, using the FALCON platform (*6*), we recently probed how diverse FFAs (i.e. environmental perturbations) induce distinct gene expression signatures in cells and how this may interact with genetic risk factors for metabolic diseases. Here, we flipped the paradigm: we probed how genome-wide perturbations (i.e. gene knockouts) influence cells’ ability to respond to lipotoxic stress. Our study serves as a resource – available to the entire scientific community – mapping hundreds of genes that influence the lipotoxicity response and the coordinated pathways that may contribute to differentiated survival. In sum, our results support further investigation into the links between altered metabolic states and organellar function – specifically how the functional status and abundance of organelles like the peroxisome shape the etiology and progression of metabolic diseases characterized by high levels of circulating lipotoxic lipids.

## Materials and Methods

### Cell culture

N1H1 immortalized kidney epithelial cells were maintained at 37°C with 5% CO2 in RenaLife Renal Basal Medium supplemented with RenaLife LifeFactors® (Lifeline Cell Technology), with the exclusion of Gentamicin and Amphotericin B. The cells were generated as outlined in Dvela-Levitt et al., 2019, with informed consent under WFUHS IRB00014033. Cells were routinely checked and were negative for mycoplasma.

### Fatty acid preparation

FFAs obtained from Nu-Chek Prep, Inc. were stored in glass vials at −20°C in the compound management facility at the Broad Institute. To prepare fatty acids, DMSO dissolved FFAs (10 µM) were added to solutions of ddH**2**O-dissolved fatty acid free BSA (Sigma #A8806) to achieve a molecular ratio of 1:6.67 BSA:FFA. Fatty acid-BSA solutions were incubated in a 37°C water bath overnight to allow for FFA-BSA conjugation. The following day, the ddH**2**O and DMSO were removed completely through full-vacuum evaporation at 37°C and centrifugation at 400g for 12h using a GeneVac HT-12 evaporator. Dried BSA-conjugated FFA crystals were resuspended with RenaLife media at room temperature for 3h on an orbital plate shaker. FFAs were filtered through 96 or 384 MultiScreenHTS HV Filter Plates (0.45 μm, Millipore, #MSHVS4510 or #MZHVN0W10) for 1 min at 500g and transferred onto cells for treatment.

### Lipidomics

#### LC-MS/MS Data Acquisition

Lipid extracts were analyzed by LC-MS/MS by General Metabolics (https://generalmetabolics.com/). 2 µL of each sample was injected and separated by UHPLC using a Nexera UHPLC system (DGU-405 degasser unit, LC40DX3 solvent delivery system, SIL-40CX3 auto sampler, CBM-40 system controller, CTO-40C column oven; Shimadzu). Separation was achieved by reverse phase liquid chromatography using an Acquity UPLC BEH C18 column (1.7 µm, 2.1 mm X 30 mm; 186002349, Waters). Separation was achieved using a 2.5 minute multi-phase linear gradient of the following buffers: Buffer A) 6:4 acetonitrile:water (v:v), 10 mM ammonium acetate, B: 9:1 isopropyl alcohol:acetonitrile(v:v), 10 mM ammonium acetate. Samples were ionized using an Optimus Turbo V + Dual TIS ion source (Sciex) and were analyzed using an X500R mass spectrometer (Sciex). Samples were analyzed in positive ionization mode and were acquired using data-dependent acquisition. MS1 data was acquired with a 0.1 second MS1 accumulation followed by top 12 MS2 data acquisition with an accumulation time of 0.03 seconds per target. Dynamic background subtraction was used and former candidate ions were excluded for 3 seconds. A collision energy of 35 V was used. With each successive measurement the exclusion list was built to include the fragmented ions from the previous run.

#### LC-MS/MS Data Processing

LC-MS/MS data processing and ion annotation was performed according to accepted protocols for mass spectrometry data processing and feature annotation. Briefly, annotation was based on matching of chromatographic retention times and MS1 values from detected features to expected retention time ranges for measured purified standards representing relevant lipid classes and/or matching of the resulting MS2 spectra from fragmentation to spectral libraries of computationally generated expected fragmentation patterns for lipids. These analyses and calculations were executed at General Metabolics, a commercial metabolomics and lipidomics service vendor.

### Proteomics

Protein extraction and tryptic digestion: 600 μg protein was reduced and alkylated at 70 °C for 15 min in lysis buffer (5 % (w/v) SDS, 100 mM TEAB pH 8.5, 40 mM CAA, 10 mM TCEP) in Low Protein Binding Microcentrifuge Tubes (Thermo Scientific, Waltham, MA, USA). The SP3 extraction protocol was followed (Hughes et al., 2019). Proteins were digested on SP3 beads in 300 µL 50 mM TEAB and 3 μg trypsin-LysC. Samples were digested overnight in a shaking incubator at 37°C. The peptide solution was separated from the beads with a magnet.

200 μg peptides were labeled with a total of 500 μg TMTpro in a final volume of 100 μL 200 mM TEAB. Unreacted TMTpro label was quenched by adding 5 μL 5% hydroxylamine in 100 mM TEAB. TMTpro-labeled samples were pooled and purified using a Sep-Pak 500 mg C18 cartridge. The C18 resin was conditioned with 80% acetonitrile in 0.2% TFA, followed by 0.2% TFA in MS-grade water. The lyophilized TMTpro pooled sample was reconstituted in 0.5% TFA and loaded onto the column. The column was then washed with 0.2% TFA and the peptides were eluted in 50% acetonitrile in 0.2% formic acid.

TMTpro pooled sample was split, and one part was prepared for a standard proteome experiment. The other part was utilized for phosphopeptide purification using the High Select TiO_2_ Phosphopeptide Enrichment kit (Thermo Fisher). The flowthrough was kept for an additional downstream enrichment using the High Select Fe-NTA Phosphopeptide Enrichment Kit (Thermo). All samples were then fractionated with the High pH Reversed-Phase Peptide Fractionation Kit (Thermo) using the following 12-step gradient of increasing acetonitrile concentrations: 5, 7.5, 10, 12.5, 15, 17.5, 20, 22.5, 25, 27.5, 30, 60%. The following fractions were then pooled together and lyophilized: 1+7, 2+8, 3+9, 4+10, 5+11, 6+12. Lyophilized fractions were resuspended in 12 μL 0.2% formic acid in MS-grade water for LC-MS analysis. The Ionopticks TS25 column was used for chromatographic separation (25 cm x 75 µm, 120 Ȧ). The peptide fractions were injected with a volume of 5 µL. Peptides were separated with a mobile phase consisting of 0.1% (v/v) formic acid in water (solution A) and 0.1% (v/v) formic acid in 80% (v/v) acetonitrile (solution B), flowing at 400 nL/min, while the column temperature was held constant at 50°C. The column underwent initial conditioning with 3% solution B, followed by a gradual linear gradient up to 40% solution B over a 120-minute period. Any remaining peptides adhering to the C18 resin were subsequently washed off with 95% solvent B for 10 minutes.

LC-MS data was collected on the VanquishNeo coupled with an Orbitrap Eclipse mass spectrometer equipped with a FAIMSpro interface, all from Thermo Fisher.

The extraction and processing of LC-Orbitrap-MS data was conducted with the PEAKS Studio 10.6 software platform following the recommendations by Schulte *et al*. (2019). The *Homo sapiens* database (Uniprot taxon ID: 9696) was used as a reference for identifying proteins using the following parameters: merge data files, parent mass error tolerance 20 ppm; fragment mass error tolerance 0.05 Da, retention time shift tolerance 5.0 min; semispecific trypsin enzyme specificity; fixed modifications: TMTpro 16plex, carbamidomethylation (C: Δm/z = 57.02); variable modifications: phosphorylation (STY: Δm/z = 79.97), oxidation (M: Δm/z = 15.99), deamidation (NQ: Δm/z = 0.98), two missed cleavages; maximal three variable posttranslational modifications per peptide). The TMT16plex reporter ions were quantified on the MS2 level with a mass error tolerance of 0.05 Da. All peptides that corresponded to false discovery rate (FDR) limits of ≤ 0.1 % and at least 1 unique identification were classified as significant protein identifications. The quantities of at least 1 peptide provided the raw protein abundances and the data was exported into .csv spreadsheets for statistical analysis. As described in (*54*) and (*55*).

Proteomics data was normalized with the trimmed mean of m-values method and differential analysis was performed with the statistical analysis package “limma.” UMAP projections of pre and post-normalized data were generated with matplotlib package. Hierarchical clustering was conducted using seaborn clustermap package using the “ward” method. KEGG pathway enrichment of clusters was determined with over-representation analysis, with a Benjamini-Hochberg procedure, and selecting pathways with FDR<25%.

### CRISPR/Cas9 knockout viability screen

Cas9-stably expressing normal kidney tubular epithelial cells were lentivirally infected with a genome-wide single guide RNA library (Broad Institute Genetic Perturbation Platform (GPP) Brunello library; Addgene, #73178) followed by puromycin selection for 48h then recovery in fresh media for 24h. FFA treatment arms included a control BSA arm and a BSA-conjugated PA treatment arm at 500 µM. At least 40 million cells were passaged per treatment arm twice a week to maintain representation of ∼80,000 constructs, averaging 500 cells infected per clone, over two weeks. Cells were harvested for genomic DNA extraction and submitted for bulk-RNA sequencing at the Broad Institute GPP. Two replicates of the screen were performed and hypergeometric distribution was used to rank sgRNAs and calculate gene p-values to generate the volcano plot. Average Log2 Fold-Change (Log2FC) was calculated as: Avg Log2FC = log2normalized reads per million (Log2RPM) PA treatment sample – Log2RPM BSA treatment sample. Genes with average -Log10(p-value)>3 were considered for subsequent analysis and follow up. Pathway analysis was performed using Gene Set Enrichment Analysis (GSEA). A pre-ranked list of all genes (ranked by -Log10(P)*SIGN(AvgLog2FC) per gene) was used to identify enriched KEGG, GOBP, and REACTOME pathways (FDR<25%).

### Gene Ontology (GO) enrichment pathway analysis

Overrepresentation of GO terms and pathway analysis was performed in R v4.3.3 using clusterProfiler v4.10.1. To set up the analysis, a common list of genes identified in both epithelial and K562 leukemia cell screens was established as a background gene set, or universe. Then, a list of differentially expressed genes (p<0.05) from each respective screen, which only included genes that were also part of the universe gene list, was used as an input vector list to identify enriched clusters. To look for enriched GO term clusters (clusterProfiler::enrichGO()), genes were converted to ENTREZ IDs, and GO type biological process (BP) was analyzed. P-values were adjusted using the Benjamini-Hochberg correction method and GO pathways with adjP<0.25 were considered enriched.

### Growth curve assay

Non-targeting control or knockout cell lines were passaged for 10 days with BSA or BSA-conjugated PA at 250 or 500 µM. Cells were passaged and counted every 3-4 days to determine the doubling rates of knockout cells compared to control cells.

### Cell line knockout generation and clonal selection

Plasmids expressing non-targeting control (NTC) (Millipore Sigma, CRISPR20-1EA), *Faf2*-targeting (5’-AGAAAGCGAAGCTCCCTTTTGG-3’, Millipore Sigma, Sanger library clone HS5000006660) *AGPS*-targeting (5’-ATAAAAATGGTAACACCTAG-3’, 5’-GTACCAATGAGTGCAAAGCG-3’) *GNPAT*-targeting (5’-ATGGCTAAAAGGCTTAACCC-3’, 5’-GTAATTCCCTTATAGACAAG-3’), *PEX3*-targeting (5’-AAATCAGAGAAATACAGGAA-3’), *PEX5*-targeting (5’-TATCTGAACTCACCAGCGAT-3’) *PEX10*-targeting (5’-CAGAAGGACGAGTACTACCG-3’), or *PEX19*-targeting (5’-AGCCACTGCGGAGTTCGAGA-3’) single guide RNAs were packaged into a second generation CMV promoter lentiviral vector system using FuGene Promega transfection agent (#E2311). Cas9-stably expressing N1H1 tubular epithelial cells were lentivirally spin-fected at 2000 RPM for 2h at 30°C with control or sgRNA containing virus with protamine sulfate (4 µg/mL). 1-2h after spinning, fresh media was added without protamine sulfate and left to incubate at 37°C overnight. Cells were selected with puromycin (4 µg/mL) for 48h followed by 24h recovery in fresh media.

For single clone selection, cells were diluted to 0.5 cells per well and plated into 96-well tissue culture plates. Cells were expanded and sequenced for mutations in the *Faf2* gene. One scrambled control clone and a clone with a mutation in FAF2 exon 6 were used for generation of FAF2-full and FAF2 domain deletion cell lines.

### Re-expression of Myc-tagged full or domain-deleted FAF2 constructs

Clonal FAF2 KO cells were virally infected with FAF2 silent mutation (5’-AGAAAGCGAAGCTCCCTTTTGG-3’ to 5’-AGGAACCGCAGTTCTCTCTTAG-3’)-containing plasmids expressing N-terminal myc-tagged full length FAF2, FAF2ΔUBX (ΔUBX=nt 3679-3927), FAF2ΔUAS (ΔUAS=nt 2974-3441), or FAF2ΔUBA (ΔUBA=nt 2644-2754) based on FAF2 NCBI transcript ID (NM_014613.3) (plasmids and sequences available upon request). Mutant constructs were packaged into a third-generation CMV promoter lentiviral vector (pLV[Exp]-Neo-CMV) and infected into Cas9-stably expressing N1H1 human kidney tubular epithelial cells at 2000 RPM for 2h at 30°C with protamine sulfate (4 µg/mL). Plasmids and corresponding virus were produced by GenScript. 1-2h after spinning, fresh media was added without protamine sulfate and left to incubate at 37°C overnight. Cells were selected with geneticin (30 µg/mL) for 72h followed by 24h recovery in fresh media.

### Viability assays

N1H1 tubular epithelial cells were seeded in PerkinElmer PhenoPlate 384-well (#6057300) or 96-well (#6055302) and maintained at 37°C with 5% CO_2_ for 48h. Seeded cells were treated with BSA or BSA-conjugated PA for 24h followed by live imaging using the Opera Phenix High Content Screening System. Hoechst 33342 (1:2000) and propidium iodide (PI) were added to cells, which were incubated for 1h at 37°C before imaging. Cell viability was determined on Harmony High-Content Imaging and Analysis Software (PerkinElmer) using Hoechst or digital phase contrast (to count number of nuclei) and PI (Thermo Fisher Scientific, #P3566) to determine the average % PI positive cells per well.

### Immunoblotting

Samples were lysed in solution containing 100 mM NaCL, 5 mM EDTA, 50 mM Tris-HCl, and 1% NP-40 (Thermo 28324) buffer, pH=7.5. Cells were incubated on ice for 20 minutes before being spun at 21,300xg for 15 min at 4°C and pellet removed. Protease (Roche, #05892791001) and phosphoprotease (Roche, #04906837001) inhibitors were added. Protein concentration was determined with Pierce BCA protein assay (Thermo #1859078). Lysed samples were prepared with 4X LDS sample buffer (Invitrogen NP0008), 10% NuPage reducing agent (Invitrogen NP0009), and to volume with ddH2O. Samples were heated to 65°C for 10min. 4-12% Bis-Tris gels (NuPage) were run at 120-150V in MOPS or MES SDS running buffer (Invitrogen #NP0002 and #NP000202). Gels were transferred to nitrocellulose membranes (BioRad #1704158) with the Trans-Blot® Turbo TM Blotting System (BioRad, #1704155) preset “Mixed MW” transfer settings. Membranes were blocked in 5% NFDM (CST 9999S) in 0.05-0.01% PBST. Primary antibodies were applied overnight in 5% NFDM at 4°C. 3X 10min washes with PBST were applied before application of secondary antibodies (1:5000). Membranes were again washed with PBST 3X by 10min and developed with Pico PLUS or Femto (Thermo #34578 and Thermo #34096). Blots were imaged with the Biorad ChemiDoc MP Touch Imaging System.

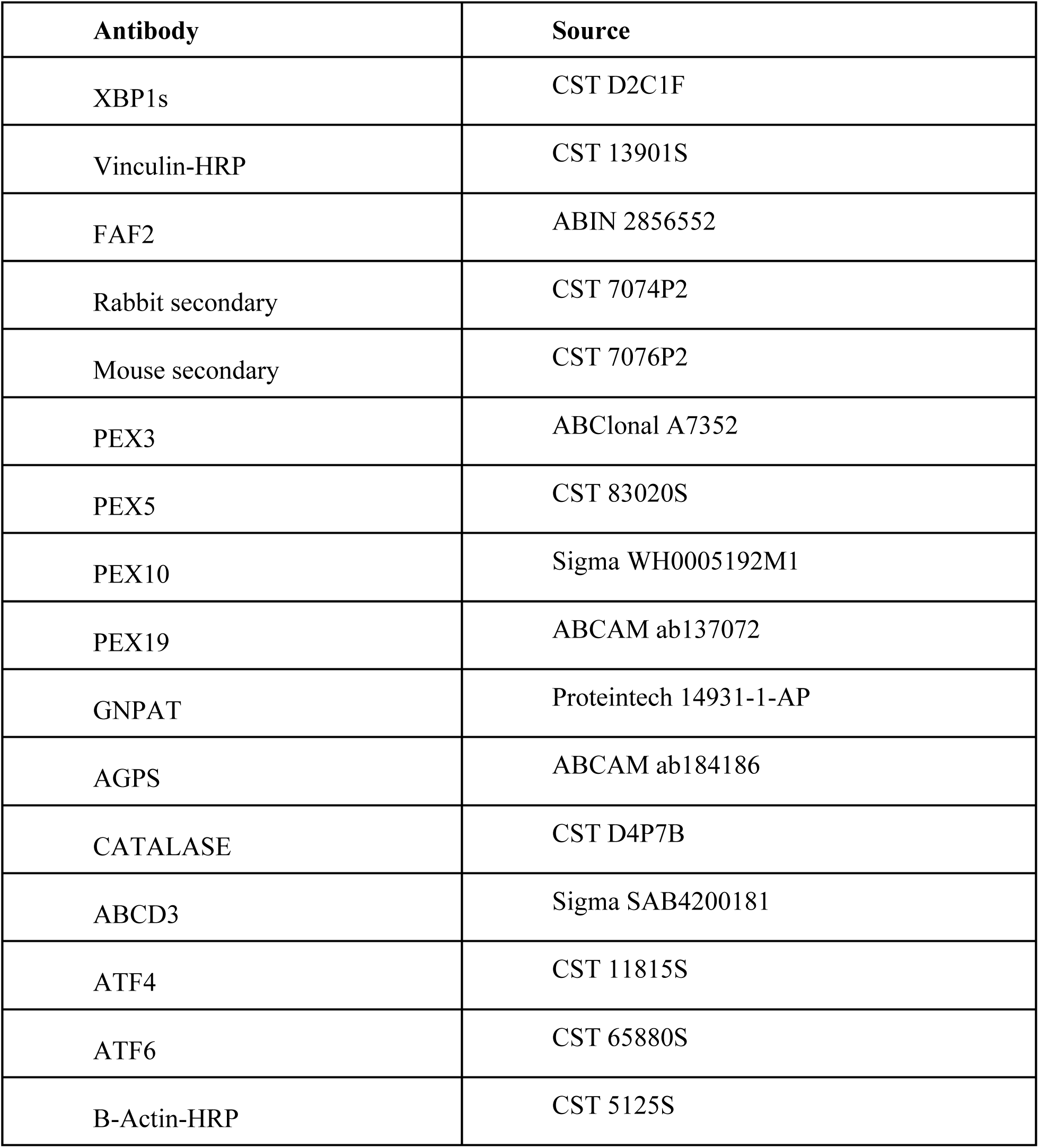

### Immunofluorescence imaging

N1H1 wildtype (WT) or KO cells were seeded at 5,000 cells per well in a PerkinElmer PhenoPlate 96-well (#6055302) and maintained at 37°C with 5% CO_2_ for 72h. Cells were fixed for 20 min in 4% paraformaldehyde (Electron Microscopy Sciences #15714), washed 2X with PBST, permeabilized for 15 min in 0.5% Triton X-100 (Sigma-Aldrich, 10789704001), washed 2X with PBST, blocked for 1h in 0.5% Roche blocking solution (Roche, #11096176001), then incubated overnight at 4°C with primary antibody diluted in blocking reagent. Cells were washed 4X with PBST, then incubated for 1h at room temperature (RT) with fluorescent-labeled secondary antibody in blocking reagent (1:500, Alexa Fluor 568 Donkey anti-Rabbit IgG (#A10042, Thermo Fisher Scientific) or anti-Mouse IgG (#A10037, Thermo Fisher Scientific)).

Nuclei were counterstained with Hoechst (1:2000, #H3570, Thermo Fisher Scientific) for 1h at RT with secondary antibody. Cells were washed 4X with PBST and imaged using the Opera Phenix High Content Screening System using a 40x or 63x water immersion objective in confocal mode. Image analysis was performed with the Harmony Imaging Software (PerkinElmer). Cell nuclei were defined using Hoechst staining, and peroxisomal puncta were quantified with intensity and spot measurements.

### Spot intensity quantification

Images were segmented into nuclei, cytoplasm, and spots after applying a Gaussian smoothing filter. Using building blocks “Calculate texture properties” and “Calculate morphology properties” the SER texture properties and STAR morphology properties were calculated respectively. Using the “Select population” building block, a linear classifier was trained on these properties using a wildtype or FAF2 knockout catalase signal. The model was trained on approximately 100 cells per genotype and then applied to the experimental groups.

### Co-immunoprecipitation (co-IP)

BSA treated N1H1 tubular epithelial cells were washed 2X with PBS before NP-40 lysis buffer was added and cells were harvested on ice. Lysates were vortexed and incubated at 4°C with rotation for 30min followed by centrifugation at 13,000g for 10min. Cell lysates were normalized by protein concentration (BCA). Lysates were incubated with 4 µg FAF2 antibody (ABIN2856552) or rabbit IgG control overnight at 4°C. 10 µL Pierce Protein A/G magnetic beads (Pierce, #88803) were added to lysates and rotated for 1h at 4°C. Supernatant was saved and beads were washed 3X in 0.05% TBS-T followed by 2x wash in PBS. IP samples were eluted with 2X NuPAGE LDS Sample Buffer (Thermo Fisher, #NP0008) with NuPAGE reducing agent (Thermo Fisher, #NP0004) in de-ionized water. Samples were loaded and run on NuPAGE Novex 4-12% Bis-Tris gels (Thermo Fisher, #WG1403BOX).

### Statistical analysis

Results are shown as mean ± SD. Visualizations and statistical significance were performed using appropriate packages in RStudio (version 1.0.143) or Graph Pad Prism 9 (two-way ANOVA with Tukey’s multiple comparisons post-hoc test unless stated otherwise in figure legends). Statistically significant differences are denoted as follows: ∗p < 0.05 ∗∗p < 0.01 ∗∗∗p < 0.001 ∗∗∗∗p < 0.0001. For all experiments, N≥3 biological replicates were examined for each condition, where one representative experiment among biological replicates is shown.

### Lipid droplet quantification

Cells were plated at a density of 8,000 cells per well in a PhenoPlate 96-well plate (PerkinElmer) and incubated for 24h at 37°C, followed by treatment with BSA-conjugated FFAs for 24h. BODIPY 505/515 (Thermo Fisher Scientific, #D3921) dissolved in 100% ethanol to 3.8 mM solution was further diluted to 3 µM then added to 2X PBS-washed cells for a 15 min incubation at 37°C. Nuclei were quantified using Hoechst 33342 nuclear stain (Thermo Fisher, #H3570). Cells were imaged using the Opera Phenix High Content Screening System and lipid droplets were quantified with Harmony High-Content Imaging and Analysis Software (PerkinElmer).

### DGAT inhibitor assay

Cells were plated at a density of 8,000 cells per well in a PhenoPlate 96 well plate (PerkinElmer) and incubated for 24h at 37°C, followed by treatment with BSA-conjugated FFAs or DGAT1 (Sigma-Aldrich, #PF-04620110) or DGAT2 (Tocris, #PF-06424439) inhibitor for 24h. Nuclei were quantified using Hoechst 33342 nuclear stain (Thermo Fisher, #H3570). Cell death was measured using propidium iodide staining (Thermo Fisher Scientific, #P3566) and lipid droplets were measured with BODIPY 505/515 dye (Thermo Fisher Scientific, #D3921). Imaging was performed using the Opera Phenix High Content Screening System and quantifications were analyzed using the Harmony High-Content Imaging and Analysis Software (PerkinElmer).

## Acknowledgments

We thank Dr. John Doench and the Broad Genetic Perturbation Platform for guidance on conducting the CRISPR screen. We thank Dr. Fabian Schulte and the Whitehead Institute’s Quantitative Proteomics core for providing consultation and experimental results. We also thank General Metabolics for providing lipidomics experimental and analysis consultations.

## Funding

National Institutes of Health grant F31 DK126252-01 (CK); National Institutes of Health grant F31 DK136312-01A1 (KRG); National Institutes of Health K00 DK123834 (MRB); National Institutes of Health (NIH) grants DK095045 and DK099465 (AG).

## Author contributions

Conceptualization: CK, KRG, AG; Methodology: CK, KRG, MRB, MKA; Investigation: CK, KRG, DB, KO; Visualization: CK, KRG; Supervision: JLP, AG; Writing–original draft: CK, KRG; Writing–review & editing: CK, KRG, JLP, AG; CK and KRG contributed equally to this work and therefore reserve the right to place their name first on their respective CVs.

## Competing interests

Authors declare that they have no competing interests.

## Data and materials availability

All data needed to evaluate the conclusions in the paper are present in the paper and/or the Supplementary Materials.

## Notes

### Competing Interest Statement

The authors have declared no competing interest.

