## Supplemental Figures S1-S5 for "FAF2 is a bifunctional regulator of peroxisomal homeostasis and saturated lipid responses"

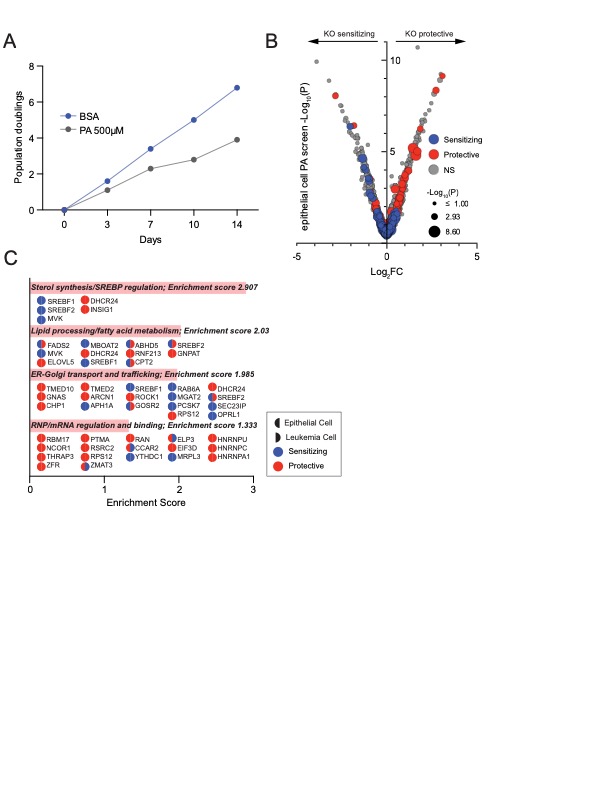

Fig. S1.

**Genome-wide screen reveals modulators of PA toxicity in diverse cell types.** (**A**) Approximate doubling times were calculated for epithelial cells treated with BSA or 500 µM PA. Cells treated with PA have reduced doubling times. (**B**) Volcano plot overlaying genome-wide screen hits from K562 human leukemia cells with the screen from epithelial cells. All genes were plotted for their Log2-fold change (Log2FC) enrichment and significance level (-Log10(P)). Color indicates sensitization (blue), protection (red), or non-significant (NS) change (grey), and size indicates level of significance from human leukemia cell screen. (**C**) Annotated genes from significantly enriched clusters in epithelial cells and K562 leukemia cells (p<0.05 or p<0.01 respectively). Enrichment scores were calculated with DAVID functional clustering analysis. The left half of each circle indicates the epithelial cell screen and the right half corresponds to the K562 leukemia cell screen. Gene knockouts that were significantly protective are in red while those that are sensitizing are in blue.

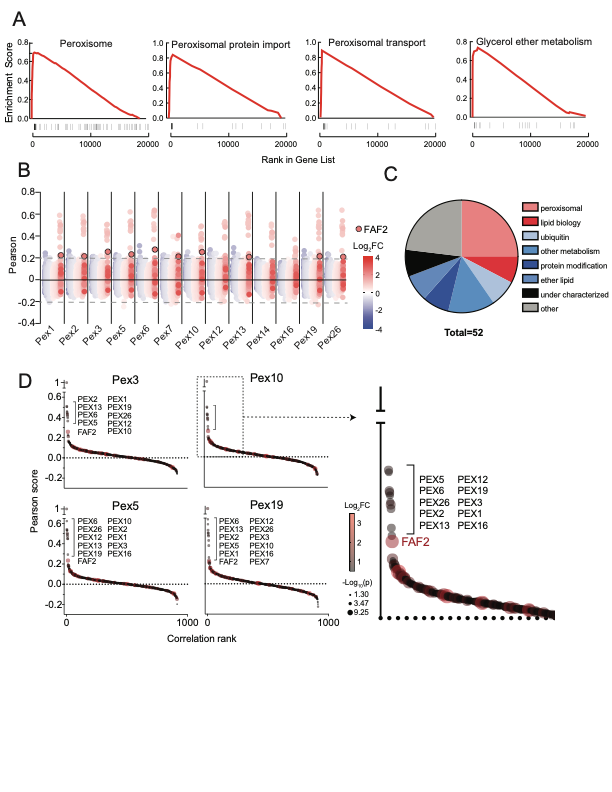

Fig. S2.

**FAF2 is a putative peroxisomal regulator.** (**A**) Representative GSEA pathways highlight protective screen hits overrepresented in peroxisomal related processes. Black lines on the x-axis indicate leading edge genes represented in the gene set list. Contribution to the enrichment score is plotted on the y-axis. (**B**) Coessentiality analysis with a manually curated gene set of peroxisomal biogenesis factors highlights FAF2 as a top emerging gene with previously unknown peroxisomal function. Pearson correlation scores were determined for all protective (red) or sensitizing (blue) screen hits against peroxisomal biogenesis factors across hundreds of cancer cell lines using DepMap CRISPR (Chronos) gene effect scores. Genes that had a Pearson score of >0.2 or <-0.2 with any PEX genes were selected and evaluated in Fig 2D and S2C. FAF2 datapoints emerged as functionally correlated with many peroxisomal biogenesis factors and are outlined in black. (**C**) Categorization of 52 genes from coessentiality analysis, many of which have primary annotated functions that are non-peroxisomal. (**D**) FAF2 is among the significantly protective hits most correlated with peroxisomal biogenesis factors (colored by Log2FC and sized according to -Log10P in the PA screen). Genes were ranked according to their Pearson correlation score.

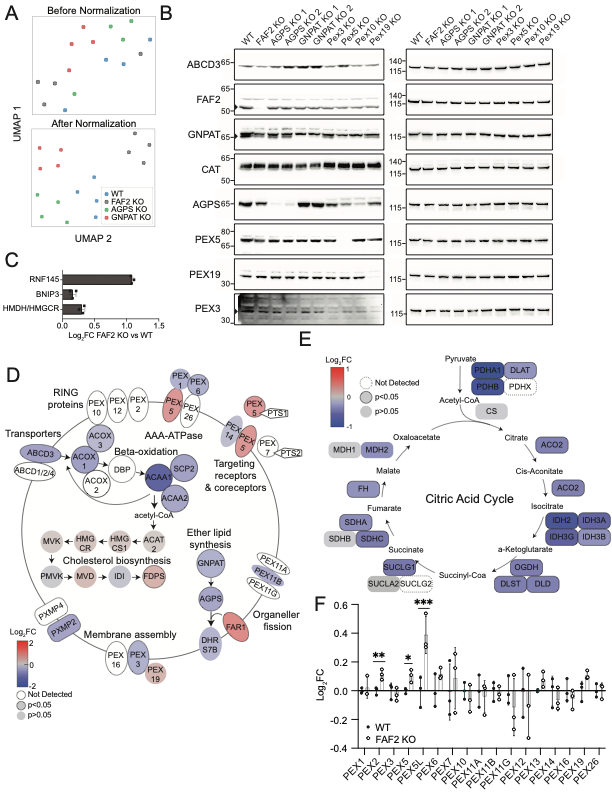

Fig. S3.

**FAF2 regulates peroxisomal protein abundance.** (**A**) Proteomic samples before and after normalization (UMAP projections). (**B**) Immunoblot of FAF2 and peroxisomal proteins in epithelial wildtype and knockout cells. (**C**) Proteomics Log2 fold change of ERAD targets of FAF2. (**D**) Representation of Log2 fold changes in peroxisomal proteins in FAF2 KO HEK cells. Genes outlined in black were significantly up or downregulated (p<0.05) in knockout cells compared to wildtype. (**E**) Log2 fold changes of mitochondrial proteins in FAF2 KO HEK cells. Genes outlined in black were significantly up or downregulated (p<0.05) in knockout cells compared to wildtype. (**F**) mRNA expression of peroxisomal gene transcripts in wildtype and FAF2 KO cells. ***p<0.001.

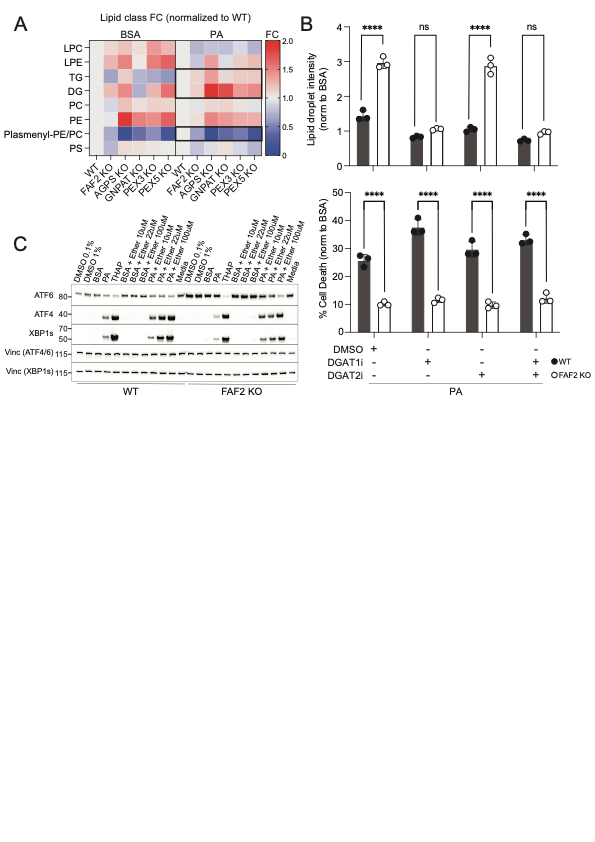

Fig. S4.

**Ether phospholipids rather than neutral lipid species are associated with FAF2 KO protection from PA.** (**A**) Heatmap of lipid species fold changes between wildtype, FAF2 KO, or peroxisomal KO cells treated with BSA or PA 350 µM shows decreased plasmenyl-PE and PCs in KO cells and increased TGs and DGs in PA-treated KO cells. (**B**) Quantification of BODIPY live imaging shows that DGAT1 inhibition (PF 04620110) but not DGAT2 inhibition (PF 06424439) prevents the increase in lipid droplet intensity in PA-treated FAF2 KO cells. PA-induced cell death measured by PI staining is not affected by DGAT1 or 2 inhibitor treatment. All PA-treated conditions were normalized to BSA. ****p<0.0001. (**C**) Western blot of ER stress markers ATF6, ATF4, and XBP1s in WT or FAF2 KO cells treated with DMSO, 100 nM Thapsigargin (THAP), or 1 mM BSA/PA with or without ether lipid for 8 h.

Fig. S5.

**Specific peroxisomal proteins are not affected by FAF2 protein levels.** Western blot showing unchanged levels of CAT, PEX5, and PEX19 protein levels in FAF2 KO or domain-deleted FAF2 mutant cells.
